# Investigating the influence of spore maturation and sporulation conditions on MalS expression during *Bacillus subtilis* spore germination

**DOI:** 10.1101/701854

**Authors:** Bhagyashree Swarge, Chahida Nafid, Norbert Vischer, Stanley Brul

## Abstract

Spore forming bacteria of the orders Bacillales and Clostridiales play a major role in food spoilage and food-borne diseases. The spores remain in a dormant state for extended periods due to their highly resistant features. When environmental conditions become favourable, they can germinate as the germinant receptors located on the spore’s inner membrane get activated via germinant binding. This leads to the formation of vegetative cells via germination and subsequent outgrowth. The present study focuses on the synthesis of protein MalS during *B. subtilis* spore germination by investigating the dynamics of the fluorescence level of a MalS-GFP fusion protein using time-lapse fluorescence microscopy. Our results show an initial increase within the first 15 minutes of germination, followed by a drop and stabilization of the fluorescence throughout the spore ripening period. Western blot analyses, however, indicate no increase in the levels of the MalS-GFP fusion protein during the first 15 minutes after the addition of the germinants. Thus the instantaneous increase in fluorescence of MalS-GFP may likely due to a change in the physical environment as the spore germination is triggered. Our findings also show that the different sporulation conditions and the maturation time of spores affect the expression of MalS-GFP and the germination behaviour of the spores.

## 1 Introduction

Sporulation and germination of genetically identical bacterial cells or spores have been extensively studied and shown to occur heterogeneously. This observation has challenged the food industries and the medical sector to extensively investigate both these processes to minimise microbiological risks by developing novel targeted strategies to eliminate spores. However, empirical studies on spores, fail to reveal the mechanistic basis of the spore’s resistance to the generally deployed bacterial elimination methods^1–3^. Therefore, gaining a detailed knowledge of sporulation and spore germination is of utmost importance. As discussed throughout this thesis, the resistance properties of spores are mainly caused by their layered structure and the chemical composition of those various layers^2,4^ and the germination receptor proteins (GRs), located in the inner membrane (IM), facilitate spore germination^5,6^. In *B. subtilis* three GRs are well known - GerA, GerB and GerK. These proteins are each composed of three subunits and can be triggered by several types of germinants including specific amino acids^4,7^. It is inferred that amino acids, such as alanine or valine, initially bind to the B-subunit of these receptors, thereby causing their activation^4,8^. Once the GRs are activated, monovalent and divalent cations are released from the spores along with the dipicolinic acid (DPA), and the degradation of the peptidoglycan cortex is ensued^7^. The latter is a prerequisite for the subsequent spore outgrowth. The occurrence of all these phases leads to complete re-hydration of the spore and restores its macromolecular synthesis and endogenous enzyme activity^6^.

During the outgrowth of spores, complex sugars, carbohydrates and organic acids can be used as carbon sources. In case of *B. subtilis*, glucose and malate are generally the preferred carbon sources^9^. In fact, any available malate is suggested to enter the TCA cycle where it can be oxidized by malic enzymes (MalS, MaeA, MleA or YtsJ) to pyruvate through oxidative decarboxylation^9^. Recently, Sinai *et al.*^10^ investigated the synthesis of a MalS-GFP reporter fusion protein over time, and predicted that MalS is one of the earliest proteins produced in the germinating *B. subtilis* spores. They observed a striking increase in the GFP intensity within the first 5 minutes of germination in spores. Remarkably, these spores remained phase-bright while the fluorescence intensity increased leading to their conclusion that protein synthesis is necessary during germination. On the contrary, a following study proved that protein synthesis is not essential for germination^11^. Parallel to these observations, in our proteomics studies ^12,13^ the relative quantity of MalS in the dormant spores is considerably high and during germination, there is no apparent change in its levels. The above-mentioned studies however, differ with respect to the age of the spores used in each study and the medium used for sporulation. Previously, it has been reported that the sporulation conditions and maturation time affect the spore characteristics and resistance properties^14^. Spores formed on solid rich media are generally more heat resistant and are characterised by a higher degree of coat proteins involved in cross-linking^14^. It has also been shown that higher spore coat protein cross-linking correlates with slight delayed germination times^11^. Therefore, with respect to MalS synthesis and MalS-GFP expression during germination the following research questions require attention: 1) Do the different sporulation conditions and spore maturation times affect MalS-GFP expression in *B. subtilis* spores? 2) Is MalS synthesized in phase-bright spores?

To answer these questions, we have compared the spores of *B. subtilis* strain PY79 and AR71 (MalS-GFP) prepared in liquid minimal medium and on solid agar medium. The germination of young (day 2) and old or mature (day 4) spores is triggered with a mixture of L-asparagine, D-glucose, D-fructose and KCl (AGFK), and with L-alanine. The time when germination starts and the actual germination time, which is the time required for the phase transition from phase-bright to phase-dark, have been analysed using time-lapse phase-contrast microscopy. The expression of MalS-GFP in young and matured spores, obtained from the liquid and solid media, is also monitored in a time dependent manner using fluorescence microscopy. In addition, the synthesis of MalS-GFP fusion protein, during the initial stages of germination, is followed by western blot analysis.

## 2 Materials & Methods

### 2.1 Bacterial strains and sporulation conditions

The *B. subtilis* wild-type strain PY79 and the mutant strain AR71 (MalS-GFP, Spc^+^; obtained from Sinai *et al.*^10^) were used in this study. Similarly, to MalS, PdhD (pyruvate dehydrogenase subunit D) was reported to be newly synthesized at an early stage during spore outgrowth^2^. Therefore, as an independent control for the fluorescence measurements, we used a *B. subtilis* strain (Erm^+^) harbouring a PdhD-IpHluorin (PC) fusion protein prepared in our laboratory. The two different sporulation media used in this study were: defined liquid medium buffered with 80mM 3-(N-morpholino) propanesulfonic acid (MOPS) and 2x Schaeffer’s medium with glucose (2x SG) agar solid medium as described previously^11,15^. Primarily, bacterial cells were cultured on a Tryptic Soy agar (TSA) plate and incubated overnight at 37°C. A single colony was inoculated in Tryptic Soy Broth (TSB) medium and incubated at 37°C at 200 rpm until the exponential growth phase was reached (OD_600_ ∼ 0.3 to 0.4). A serial dilution was subsequently performed of the culture in MOPS liquid medium or 2x SG liquid medium and incubated overnight at 37°C while shaking at 200 rpm. A single dilution containing exponentially growing cells was selected and the bacterial culture was further enriched in 20 ml MOPS liquid medium or 2xSG liquid medium. For sporulation, cells cultured in MOPS liquid medium were inoculated into 300 ml MOPS liquid medium and cultured at 37°C while shaking at 200 rpm. In contrast, cells cultured in 2x SG liquid medium were spread on 10 plates containing 2x SG agar and incubated at 37°C.

### 2.2 Spore harvesting

Spores cultured in MOPS liquid medium and on 2x SG solid medium were harvested after 48 and 96 hrs^14^. Spore samples were extensively washed with pre-chilled milliQ-water at 4°C. Subsequently, Tween-20 (0.01% v/v) was added to the spores in milliQ-water to kill vegetative cells and to improve purification. The harvested spores were examined under the microscope using a haemocytometer and the spore yield was estimated. Remaining vegetative cells and/or germinated cells (phase dark) were removed using the Histodenz gradient centrifugation method describe by Abhyankar *et al.^14^*. The purified harvested spores contained more than 99% phase bright spores. Spores were stored at −80°C until further use.

### 2.3 Germination and Fluorescence measurements

For time-lapse germination experiments, spores were heat activated at 70°C for 30 minutes at 100 rpm and placed on ice for 10 min. Immediately after incubation on ice, the spores were transferred to an agarose pad containing 2% w/v agarose mixed with a double concentration of MOPS medium and germinants (AGFK and L-alanine) in a 1:1 ratio. The set-up of the microscope slide for time-lapse microscopy imaging has been described by Pandey *et al.*^16^. Time-lapse microscopy imaging was performed at 37°C in a time dependant manner for 2 hours using a temperature-controlled incubation system. A Nikon ECLIPSE Ti-E inverted widefield fluorescence time-lapse microscope equipped with an Apo TIRF 100x H/1.49 objective and a Hamamatsu ORCA-AG cooled charge-coupled device (CCD) camera with integrated Perfect Focus System was used. Phase contrast images were taken with an acquisition time of 100 ms. Fluorescence images were taken with a GFP filter and IpHluorin filter with an acquisition time of 250 ms and at 50% power. Phase-contrast and GFP-fluorescence time-lapse series were recorded at a sample frequency of 4 to 7 frames per 5 min, with the requirements of about 100 recorded spores per measurement. One replicate for control PdhD-IpHluorin strain was performed. Spore germination times and fluorescence intensity were both analyzed using SporeTrackerX^17^(https://sils.fnwi.uva.nl/bcb/objectj/examples/sporetrackerx/MD/sporetrackerx.html), a plugin for ObjectJ (https://sils.fnwi.uva.nl/bcb/objectj/). The time required for initiating the phase bright to phase dark transition after addition of germinants (henceforth referred to as Start of Germination) and time required to complete the phase transition (referred to as Germination time) were analysed. For GFP measurements, the intensity of a wild-type strain, lacking the *gfp* gene, was subtracted from the net average fluorescence intensity. All statistical analysis was performed in SigmaPlot 13.0 (Systat Software Inc.). To analyse results of live-imaging data from two independent replicates student’s *t*-test was performed and p-values determined.

### 2.4 Western Blot analysis

Spores of *B. subtilis* mutant strain AR71 (MalS-GFP), prepared on solid and liquid media, were heat activated at 70°C for 30 min. Germination was triggered using L-alanine and an AGFK mixture. Spores were harvested before (t = 0 min) and after the addition of the germinants (t = 5, 15 min). To stop further protein synthesis during sample processing, chloramphenicol (100 µg/ml) and methanol (20% m/v) were added to the samples from each time point. The spores were mixed in phosphate buffered saline (PBS) containing protease inhibitor cocktail (cOmplete™, Mini, EDTA-free Protease Inhibitor Cocktail, Roche) and lysed using a Precellys 24 homogenizer (Bertin Technologies, Aix en Provence, France, 6.5, 40s, 7 cycles). The protein concentration of each protein sample was estimated using a Bradford assay kit (Bio-rad, Utrecht). For every time point, equal concentration of proteins were incubated at 100°C for 10 min with Laemmli sample buffer. The proteins were separated on a 10% Bis-Tris SDS gel and electro blotted overnight onto a polyvinylidene difluoride (PVDF) transfer membrane. The PageRuler™ Prestained Protein Ladder, 10 to 180 kDa, was used as a reference ladder. For Immunoblot analysis of GFP fusion proteins, the membranes were blocked for 1 hr at room temperature with a mixture containing 0.05% Tween-20, 5% skim milk in 1X Tris buffered saline (TBS), pH 7.4. Blots were then incubated for 1 hr at room temperature with polyclonal rabbit anti-GFP antibodies (1:10,000 in 0.05% Tween-20, 5% skim milk in 1X TBS pH 7.4). Subsequently, the membranes were incubated for overnight at 4°C with peroxidase conjugated anti-rabbit secondary antibody (1:10,000 in 0.05% Tween-20, 5% skim milk in 1X TBS pH 7.4). The SuperSignal™ West Femto Maximum Sensitivity Substrate kit was used for final detection of the protein on the blot.

## 3 Results

### 3.1 Determining the effect of maturation and sporulation conditions on the germination behaviour of spores of B. subtilis strain AR71 (MalS-GFP) and the wild type strain PY79

All the young (day 2) spores of stain AR71 prepared in MOPS liquid medium differ significantly from their matured (day 4) partners with respect to the time required to start germination. At the population level, the relatively young spores are less prone to initiate germination in comparison to the older spores (45% young spores vs. 61% matured spores in 10 min; Table 1(A)). However, these two spore populations do not differ significantly in their germination time (Figure 1, Table 1(A)). The time required to start germination and the germination time differ significantly in case of the young and mature spores of the wild type strain PY79 (Table 1 (B), Figure 1). Furthermore, the young wild-type spores initiate germination later than the matured spores (35% vs. 44% in 10 min, Table 1 (B)).

**Table 1.**
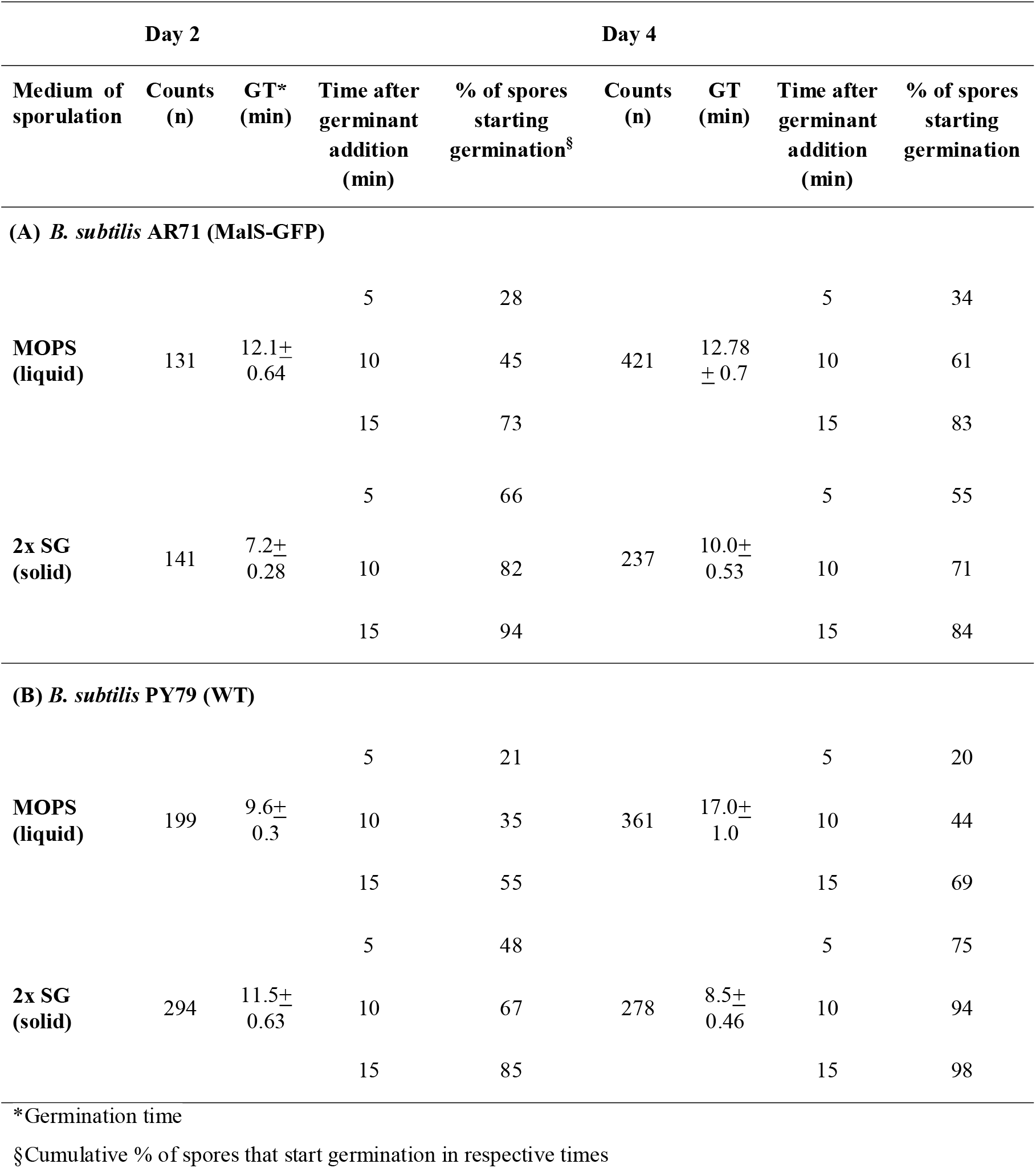
Germination behaviour of young and mature spores of *B. subtilis* wild type strain PY79 and mutant strain AR71 (MalS-GFP) prepared in liquid (MOPS) and solid (2x SG) media.

| Day 2 |  |  |  |  | Day 4 |  |  |  |
| --- | --- | --- | --- | --- | --- | --- | --- | --- |
| Medium of sporulation | Counts (n) | GT* (min) | Time after germinant addition (min) | % of spores starting germination<sup>§</sup> | Counts (n) | GT (min) | Time after germinant addition (min) | % of spores starting germination |
| (A) *B. subtilis* AR71 (MalS-GFP) |  |  |  |  |  |  |  |  |
| MOPS (liquid) | 131 | 12.1±0.64 | 5 | 28 | 421 | 12.78±0.7 | 5 | 34 |
|  |  |  | 10 | 45 |  |  | 10 | 61 |
|  |  |  | 15 | 73 |  |  | 15 | 83 |
| 2x SG (solid) | 141 | 7.2±0.28 | 5 | 66 | 237 | 10.0±0.53 | 5 | 55 |
|  |  |  | 10 | 82 |  |  | 10 | 71 |
|  |  |  | 15 | 94 |  |  | 15 | 84 |
| (B) *B. subtilis* PY79 (WT) |  |  |  |  |  |  |  |  |
| MOPS (liquid) | 199 | 9.6±0.3 | 5 | 21 | 361 | 17.0±1.0 | 5 | 20 |
|  |  |  | 10 | 35 |  |  | 10 | 44 |
|  |  |  | 15 | 55 |  |  | 15 | 69 |
| 2x SG (solid) | 294 | 11.5±0.63 | 5 | 48 | 278 | 8.5±0.46 | 5 | 75 |
|  |  |  | 10 | 67 |  |  | 10 | 94 |
|  |  |  | 15 | 85 |  |  | 15 | 98 |

\*Germination time
§Cumulative % of spores that start germination in respective times

**Figure 1.**
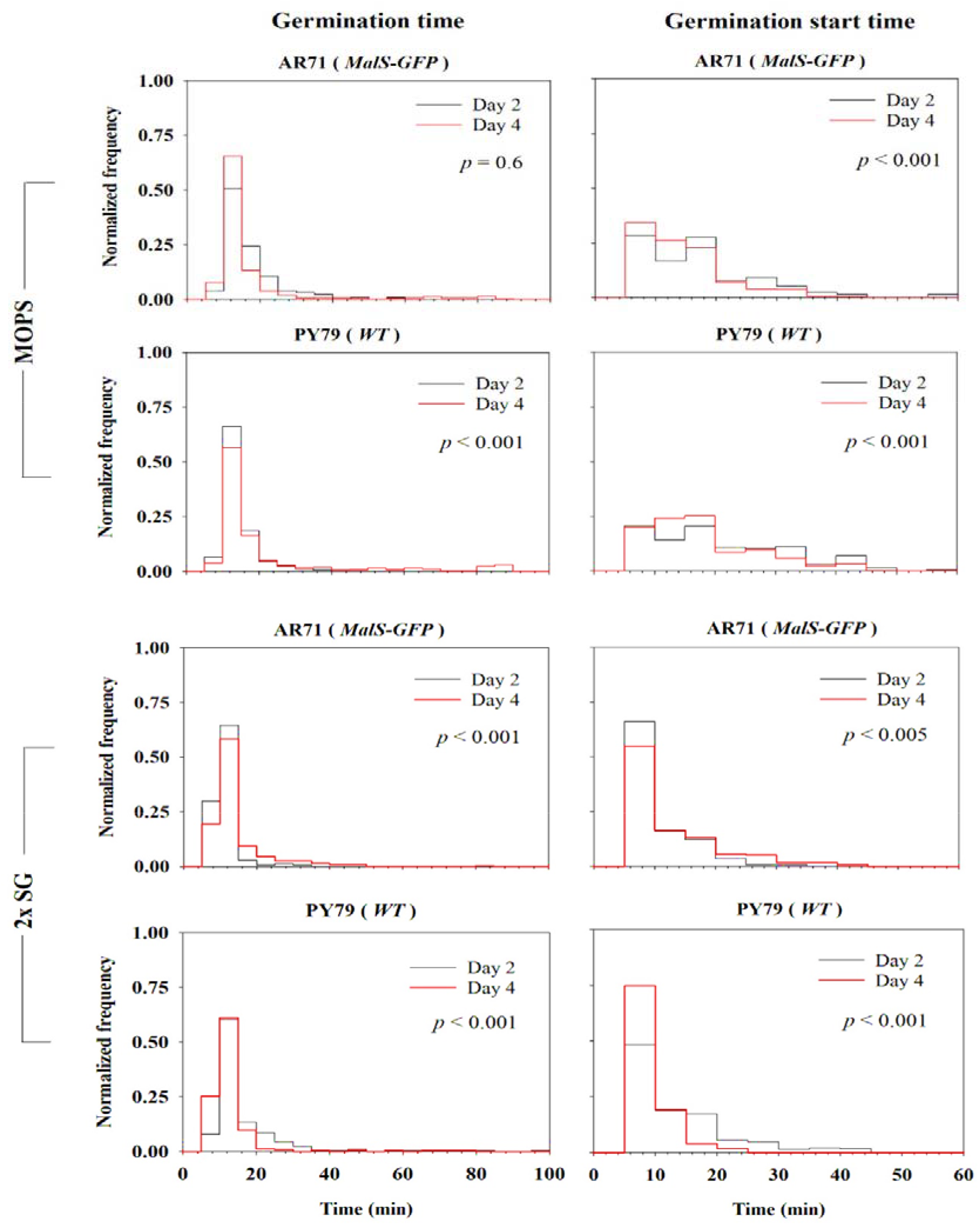
Effect of maturation time on germination behaviour. Frequency distribution curves of *B. subtilis* wild type strain PY79 and strain AR71 spores prepared in MOPS and 2x SG media. Germination time and start of germination are illustrated. Young spores (black line) are overlaid with mature spores (red line). The significance of the data is tested using the student’s *t*-test. The Histogram is normalized based on the total spore count. Observations of two biological replicates are grouped and analysed as one data set.

In case of the sporulation on 2x SG solid medium, the younger AR71 spores are more prone to start germination than their matured counterparts (82% vs 71% in 10 min, Table 1(A)). On the contrary, the younger PY79 spores are less prone to start germination compared to the mature spores (67% vs. 94% in 10 min, Table 1 (B)). In general, with the exception of AR71 old spores, the wild type PY79 and mutant strain AR71 spores prepared on the solid medium start germination earlier than those prepared in a liquid medium. (Figure 2, Table 1).

**Figure 2.**
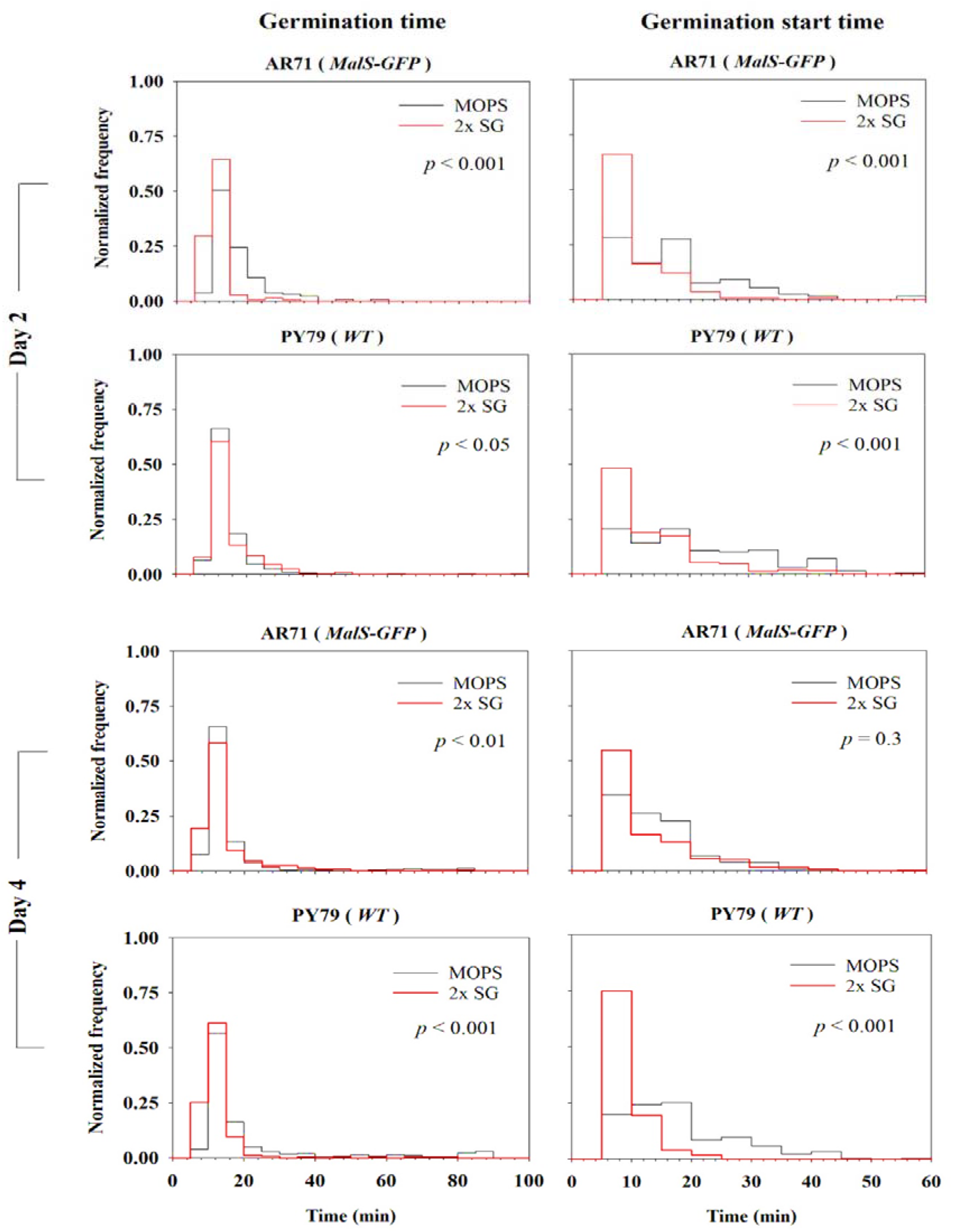
Effect of the sporulation condition on the germination behaviour of young and matured spores of wild type strain PY79 (WT) and strain AR71 (MalS-GFP). The histogram is normalized based on the total spore count. Observations of two biological replicates are grouped and analysed as one data set. The significance of the data is determined using the student’s *t*-test.

### 3.2 Determining the effect of the maturation time and sporulation conditions on MalS-GFP fluorescence

The single spore imaging results obtained for *B. subtilis* AR71 (MalS-GFP) spores during spore germination and outgrowth show an initial increase in MalS-GFP fluorescence signal for both the spore batches prepared in liquid and on solid media. After approximately 30 min the fluorescence signal decreases and stabilizes throughout the spore ripening period in spores produced in both culturing conditions. In contrast, the fluorescence intensity of PdhD-IpHluorin decreases within the first 30 min of germination and stabilizes for the rest of the measurements (**Supplementary figure 2**). The collage of the individual representative spores, of differing the culture condition and maturation times, shown in Figure 3 depicts the changing localization and distribution of the GFP fluorescence over time. The young phase-bright MalS-GFP spores, sporulated in both liquid and on solid medium, show a clustered GFP fluorescence. Upon completion of germination, the clustered GFP diffuses throughout the spore. However, in some of the control (PdhD-IpHluorin) spores (day 2, MOPS) the GFP clusters are still present even after germination is completed (**Supplementary Figure 2**).

**Figure 3.**
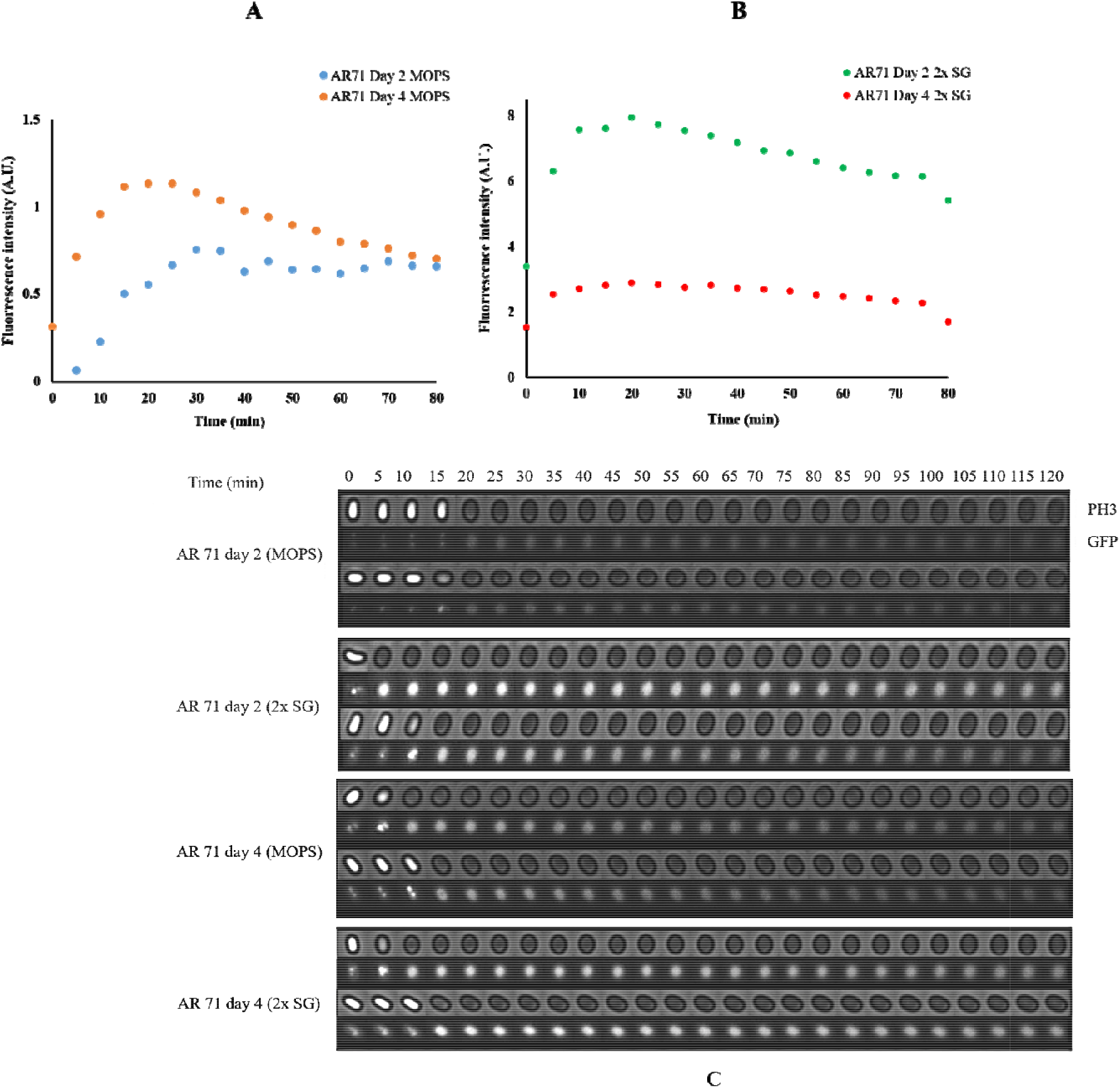
Real-time expression of MalS-GFP during spore germination. **(A) & (B) The changes in fluorescence intensity of MalS-GFP over time.** The fluorescence intensity of spores from germinating day 2 and day 4 spores, cultured in MOPS liquid medium and in 2x SG solid medium, are measured over a period of 80 min. The fluorescence intensity values of the wild type spores PY79 are subtracted from the measured fluorescence values of MalS-GFP to eliminate the background fluorescence of the spore itself. **(C) A collage of the images obtained of the *B. subtilis* AR71 (MalS-GFP) young and mature spores, prepared on solid and liquid media, while germinating.** Images of each individual spore is captured every 5 min for a duration of 2 hrs to observe the phase transition (phase-bright to phase-dark) and the changes in fluorescence intensity of MalS-GFP. As representative for the data-set only two spores per strain are shown here.

**Figure 4.**
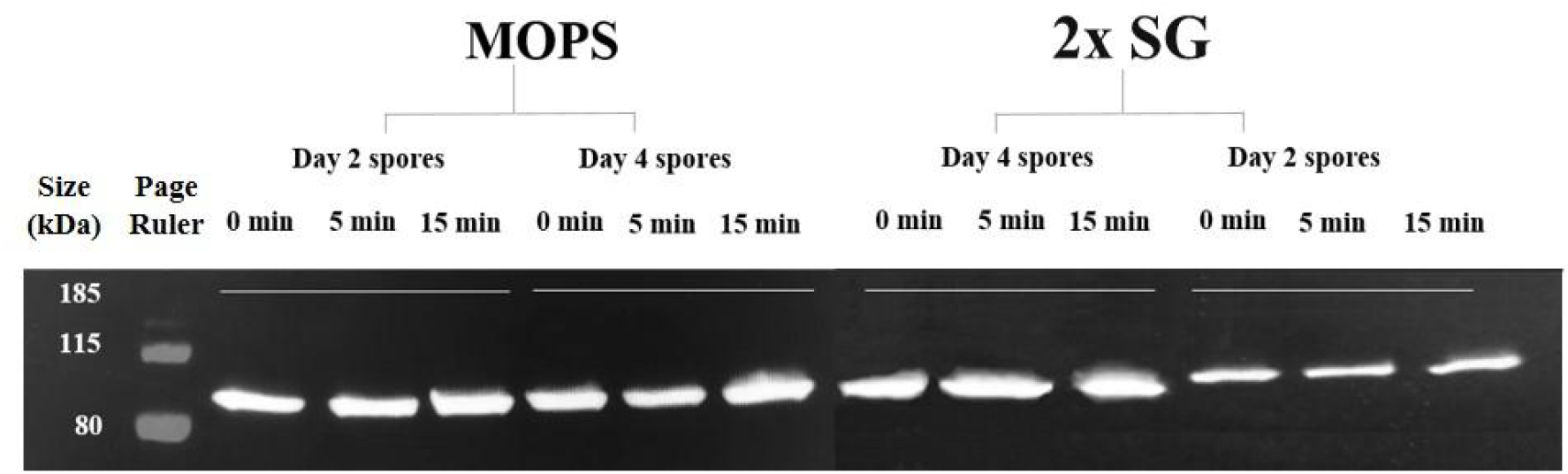
Western blot analysis of MalS-GFP fusion protein. Dormant spores are induced to germinate by various germinants (see **Materials and Methods**). Protein is extracted from dormant spores harvested before (t = 0) and after germinant addition (t= 5, 15 min). For day 2 (young) spores prepared on 2x SG 15 µg of protein whereas for day 4 (old) spores 30 µg of protein was loaded for each time point. For day 2 (young) and day 4 (old) spores prepared in MOPS 30 µg protein for each time point loaded on the gel respectively. Spores from liquid medium are phase bright at t = 0 and 5 min and completed the germination at 15 min. Spores obtained from solid medium completed the germination at t = 5 min. Protein extraction is carried out as shown by Sinai *et al.*^10^

### 3.4 Effect of sporulation conditions and maturation time on the levels of MalS-GFP fusion protein

The western blot analyses show that in young and mature AR71 spores, prepared in liquid and on solid media, the levels of the MalS-GFP fusion protein do not increase significantly within 15 minutes after the addition of the germinants. The microscopy images show that by this point all spores turn phase dark i.e. their germination is completed.

## 4 Discussion

Dormant spores of *Bacillus subtilis* return to life through germination in a sequential order of steps. During the first steps of revival, the dormant spores release cations, small molecules, and CaDPA from the core. This triggers cortex hydrolysis, further CaDPA release and full core hydration. Completion of germination results in a phase-dark spore which is ready to start macromolecular synthesis. The germination process itself is very heterogeneous and germination kinetics appears to be affected by different sporulation conditions^18^ and spore maturation^11^. Also, the recent evidence shows that the spore-coat characteristics differ in the spores prepared in the liquid and on solid media. Nevertheless, the precise effect of spore maturation and sporulation conditions on germination remains unclear. Evidences over decades of work on spore germination indicate that there is no significant metabolic activity and no protein synthesis in a spore until completion of germination^4–6,19^. Albeit, a recent report has shown protein synthesis to be necessary during germination, and dormant spores to be metabolically active^10^. The discrepancy between the earlier studies and the work of Sinai and colleagues may stem from the differences in the sporulation conditions and age of spores used in the respective studies. Therefore, in the present work we attempt to observe the effect of sporulation conditions and spore maturation on behaviour of spore and protein synthesis during germination.

With regards to the germination behaviour, we focus on two different aspects: the effect on (a) the time required start germination and (b) the actual germination time (total duration to achieve phase darkening). In general, the spores prepared on solid medium start germination earlier as compared to spores sporulated in the liquid medium. This is in accordance with a previous report^18^. Moreover, the young AR71 (MalS-GFP) spores, prepared in both liquid and solid media, complete germination faster than the matured spores. Interestingly, the young wild type spores prepared in liquid medium behave similarly by having a shorter germination time than that of the matured spores, but the spores prepared on the solid medium differ in this aspect. Thus, for the wild type spores prepared on solid medium the age or the maturation time of spores appears to affect the germination time as well. This could be due to the difference in (a) thickness of the spore-coat layers and the protein-crosslinking therein^14^ or (b) the number of germination related proteins in the spores, as suggested previously^8^.

Next, we analyse the influence of maturation times and sporulation conditions on MalS-GFP expression over time. Interestingly, for the spores prepared on solid medium the MalS-GFP fluorescence is always higher than that for spores prepared in liquid medium. This distinction might be a variation in GFP maturation behaviour caused by the differing growth conditions, as suggested previously^20^. Furthermore, MalS-GFP in the spores appears to be clustered until germination is completed. Previously, it has been shown that a pH of 6.5 promotes aggregation of EGFP^21^. Since the pH in the spore core is between 6.0 and 6.4^22,23^, the observed clustering of MalS-GFP could represent GFP aggregation. The observation that these clusters diffuse throughout the spores during the ripening stage can be attributed to the phase transition in the state of core water as a result of initial heat activation^24^. The decrease in MalS-GFP fluorescence after 30 min may also result from the changed state of core water, the water intake by a germinating spore or due to utilization of MalS by the reviving spore. However, our proteomics and transcriptomic data^13^ do not support the MalS utilization theory.

The western blot analyses show no variation in the MalS-GFP levels between the germinated and dormant spores. Yet, the MalS-GFP fluorescence of both the young and matured spores, prepared in solid and liquid medium, increases during the initial 15 min after the addition of the germinants and then stabilizes after a decline. On the contrary, for the control *B. subtilis* PdhD-IpHluorin spores, the GFP fluorescence gradually decreases within 15 min. At this stage the spores have transitioned from the phase-bright into the phase-dark state i.e. they have completed germination (**Supplementary figure 1**). The increase in MalS fluorescence intensity may be linked with proper folding of the GFP or its maturation. GFP maturation requires molecular oxygen^25^. Interestingly, in *Bacillus megaterium* spores previous experimental work has shown that heat activation increases the respiration rate and thereby likely promotes the occurrence of higher molecular oxygen levels in spores^26^. Similarly, in the heat activated AR71 spores, the molecular oxygen may become available during the initial germination stages and might dictate maturation of any pre-synthesized MalS-GFP accumulated during sporulation. Alternatively, a dehydrated intermediate of GFP has been identified^27^, which requires a single protonation and dehydration cycle to attain functional maturity. This process can be carried out when spores initiate the transfer of protons during germination as the spore core rehydrates. For further characterization and verification of the functionality of the GFP label in spores, it is recommended to produce and express the GFP in the spore core without having it fused onto any specific protein. Based on our results and given the fact that the maturation times for a variety of GFPs range between 4 to 28 min^28^, a *de novo* synthesis of MalS-GFP fusion followed by its folding within 5 min after the addition of the germinant, as suggested by Sinai *et al*^10^ appears to be conceptually challenging.

In conclusion, our study shows that the sporulation conditions and maturation period of spores affect their germination behaviour and the fluorescence intensities whereas expression of MalS-GFP and thereby protein synthesis in germinating spore remain unaffected.

## Notes

#### Summary of Updates

The author name Norbert Fischer is corrected to Norbert Vischer.

