## Supplementary Figures for "Investigating the influence of spore maturation and sporulation conditions on MalS expression during *Bacillus subtilis* spore germination" (from another version of this preprint: version 1, 2019-07-13)

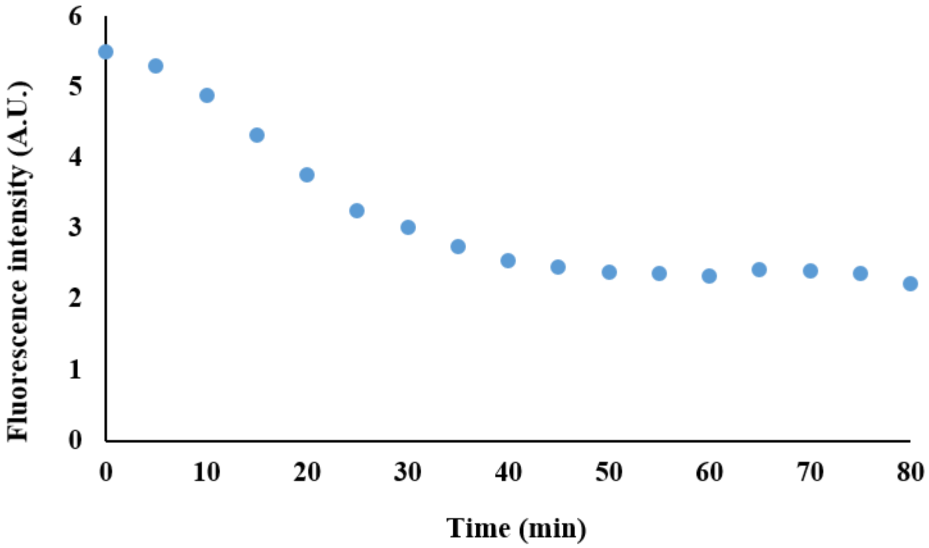


**Supplementary Figure 1: Monitoring the change in fluorescence intensity of young *B. subtilis* strain PdhD- IpHluorin spores (MOPS) during germination**

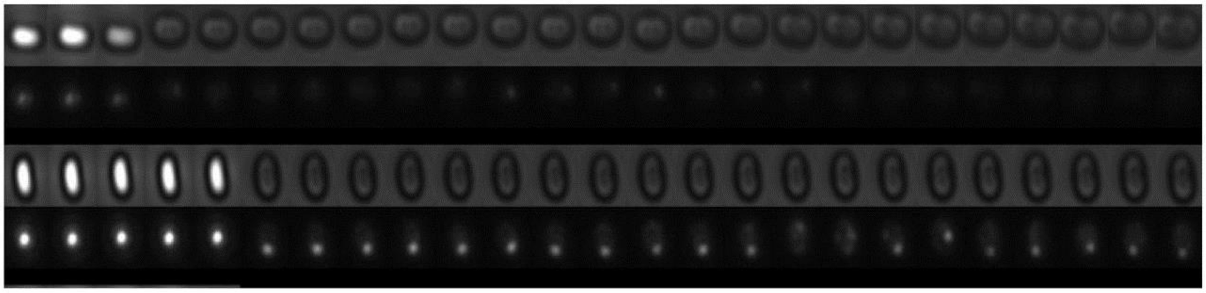

**Supplementary Figure 2: Germination over time of young *B. subtilis* PdhD-IpHluorin spores grown in MOPS medium.** Shown is a collage of germinating single spores that are captured on different time points. Two individual spores per strain are shown. Images of each individual spore are captured every 5 min using two different channels: PH3 (phase contrast) and GFP. The PH3 channel visualizes the germination over time (phase bright to phase dark). The GFP channel visualizes the increase in fluorescence intensity over time. No control is done with PdhD D4 spores.
